# Electromechanical Dynamics and Myogenic Responses in Cerebral Smooth Muscle Cells and Capillary Pericytes

**DOI:** 10.64898/2026.04.03.715998

**Authors:** Niloufar Khakpour, Maria Sancho, Nicholas R. Klug, Hannah R. Ferris, Fabrice Dabertrand, Mark T. Nelson, Nikolaos M. Tsoukias

## Abstract

Cerebral blood flow (CBF) control is essential for normal brain function and is disrupted in pathological conditions. Arterial diameters are tightly regulated to provide on demand increases in blood flow in regions of neuronal activity. Pericytes (PCs) exhibit robust myogenic tone and may also respond to neuronal activity to fine-tune local resistance and blood flow. Thus, mural control of microcirculatory resistance may extend beyond arteries and arterioles. Yet, PC’s electrophysiology and contractility have not been thoroughly characterized, and this prohibits an integrated view of brain blood flow control. In this study, we develop a detailed mathematical model of mural cell electrophysiology, Ca^2+^ dynamics and biomechanics. The model is informed by electrophysiological data in smooth muscle cells (SMCs) or PCs and predictions are compared against pressure-induced responses in isolated arterioles and capillaries, respectively. Simulations recapitulate myogenic constrictions and examine differences in contractile dynamics as we move from arterioles to proximal and distal capillaries. In arteriole-to-capillary transitional (ACT) zone PCs, increased mechanosensitivity, more Ca^2+^ influx through non-selective cation (NSC) channels and/or a higher sensitivity of the contractile apparatus to Ca^2+^ can compensate for reduced L-type voltage-operated (VOCC) Ca^2+^ influx and allow for robust constrictions at the lower operating pressures of capillaries relative to the arterioles. A significant Ca^2+^ influx through NSC relative to VOCC, however, can decouple the PC’s contractile apparatus from electrical signaling. Vasoactivity to chemomechanical stimuli along the arteriole to capillary axis is progressively driven by VOCC-independent Ca^2+^ influx and Ca^2+^ sensitization with slow kinetics. The proposed cell model can form the basis for detailed multiscale and multicellular models that will examine physiological function at a single vessel or vascular network levels and investigate CBF control in health and in disease.

**Key points:**

- A mural cell model of electrophysiology, calcium (Ca^2+^) dynamics and biomechanics is informed by data and adapted for modeling cerebral arteriole smooth muscle cells and capillary pericytes.
- Ion channel activities are characterized by patch-clamp electrophysiology in isolated cerebral smooth muscle cell and pericytes, and capillary and arteriole electromechanical responses to transmural pressure changes are assessed using novel ex vivo preparations.
- Myogenic constrictions in arterioles can be reproduced by pressure-induced non-selective cation channel (NSC) activation that depolarizes the cell, opens L-type Ca^2+^ channels (VOCCs) and increases Ca^2+^ influx.
- Robust myogenic constrictions in arteriole-to-capillary transition (ACT) zone pericytes may reflect significant Ca^2+^ influx through NSC, increased mechanosensitivity, or higher sensitivity of the contractile apparatus to Ca^2+^, potentially compensating for reduced VOCC density relative to arteriolar smooth muscle.
- A significant contribution of NSC relative to VOCC in Ca^2+^ influx, can decouple the contractile apparatus from electrical signaling.
- The model shows how gradients in ionic activities, mechanosensitivity and/or Ca^2+^ sensitivity can alter contractile phenotype and electromechanical coupling along the arteriole to capillary continuum.
- The proposed model can form the basis for detailed multiscale and multicellular models that will investigate cerebral blood flow control in health and in disease.

## Introduction

Blood is delivered to the brain through pial surface vessels that supply penetrating/parenchymal arterioles (PAs) and a dense capillary network. Capillaries are strategically positioned near every neuron, ensuring efficient delivery of oxygen and nutrients and the removal of waste byproducts. The brain is an organ of high metabolic activity that resides within the limited space of the skull. It relies on a tightly regulated microvascular network to continuously match supply and demand at a local level, while maintaining essentially constant total perfusion and blood volume at the organ level. Microvascular diameters are dynamically regulated via autoregulatory mechanisms and metabolic feedback, while feedforward control, through neurovascular communication, allows rapid on-demand increases in blood flow at regions of neuronal activity. Proper vascular control is essential for maintaining normal brain function and is disrupted in pathological conditions such as Alzheimer’s disease and stroke (Zlokovic, 2011; Kisler *et al*., 2017*a*; Anderle *et al*., 2025; Santisteban & Iadecola, 2025). How the microvascular network maintains appropriate perfusion in the presence of steep spatiotemporal gradients in pressure, lumen diameter, or metabolic demands remains incompletely understood.

Within this hierarchical network, microvascular mural cells serve as the key modulators of cerebral blood flow (CBF). Smooth muscle cells (SMCs) encircling arteries and arterioles act as sensors of vasoactive signals and as actuators to adjust vessel diameter and perfusion. They exhibit robust myogenic responses to pressure changes and respond rapidly to nearby neuronal activity for on demand local increases in CBF (Knot & Nelson, 1998; Davis & Hill, 1999; Attwell *et al*., 2010). Their contractile responses to chemo-mechanical stimuli are associated with depolarization and Ca^2+^ influx to engage the contractile apparatus; arteriolar tone is tightly coupled to membrane potential (V_m_) and free intracellular calcium ([Ca^2+^]_i_) dynamics.

Downstream, pericytes (PCs) enwrap cerebral capillaries. Their density and morphology changes along the capillary network with ensheathing PCs covering proximal to the arteriole capillaries while thin strand PCs more sparsely populate distal capillaries (Hartmann *et al*., 2015; Grant *et al*., 2019). PCs were considered passive structural elements of the cerebrovascular wall. However, arteriole-to-capillary transitional (ACT) zone (first 3-4 vessel generations proximal to the arteriole) PCs exhibit myogenic constriction to pressure elevations like SMCs (Klug *et al*., 2023; Ferris *et al*., 2025) while PCs along distal capillary segments may constrict with slower kinetics (Hartmann *et al*., 2021; Klug *et al*., 2023). PCs may fine-tune local resistance and mediate neurovascular coupling (Hamilton *et al*., 2010; Hall *et al*., 2014; Hartmann *et al*., 2015), while their dysfunction may contribute to perfusion deficits in brain pathologies (Kisler *et al*., 2017*b*; Stamenkovic *et al*., 2025). Thus, the emerging view is that SMCs and PCs form a continuum of mural control extending from arteries to distal capillaries. However, the distinct electrophysiological and mechanical behaviors of the two cell types are not well characterized nor how their modulation of arteriole- and capillary-tone intersects to coordinate local perfusion.

Decades of computational modeling of V_m_ and Ca^2+^ dynamics in excitable cells including cardiac cells or neurons have advanced physiological investigations in the heart or the brain. This progress has not been paralleled in the vasculature and there is a scarcity of similar models for vascular mural or endothelial cells. We have previously reviewed early attempts in modeling vascular SMC electrophysiology (Tsoukias, 2011). Some of the first attempts to model SMC Ca^2+^ dynamics include works by Wong and Klassen (Wong & Klassen, 1993), Gonzalez-Fernandez and Ermentrout (Gonzalez-Fernandez & Ermentrout, 1994), Fink and coworkers (Fink *et al*., 1999) and Parthimos and coworkers (Parthimos *et al*., 1999). However, the first detailed SMC model that used experimentally characterized ionic currents was presented by Yang et al. (Yang *et al*., 2003) for a cerebral artery SMC. The model integrated electrochemical dynamics with biomechanics and formed the basis or inspired subsequent efforts by Edwards and Pallone (Edwards & Pallone, 2007), Jacobsen and coworkers (Jacobsen *et al*., 2007), Karlin (Karlin, 2015), or our group (Kapela *et al*., 2008). More resent works focused on utilizing adaptations of these earlier models into multicellular arrangements or coupling with other cell types to investigate aspects of intercellular communication and signaling or integrated cell and vessel function (Koenigsberger *et al*., 2006; Christian Brings Jacobsen *et al*., 2007; Kapela *et al*., 2010*a*; Dormanns *et al*., 2015*b*, 2015*a*). Progress in the field has been limited by the difficulty in characterizing weak transmembrane currents present in mural cells and the significant variability between mural cells from different vascular beds or from different size vessels within the same vascular bed.

In this study we utilize experimental patch-clamp data to characterize main transmembrane currents in freshly isolated capillary PCs. We compare current densities with those of cerebral SMCs and explore the physiological relevance of these differences for integrated cell/vessel responses using mathematical modeling. Based on this data, we propose working models of electrophysiology, Ca^2+^ dynamics and biomechanics in cerebral PCs and SMCs, capable of recapitulating myogenic responses in brain arterioles and capillaries. Ex vivo measurements of capillary and arteriole responses to pressure elevations provide key constraints for these models. Finally, we provide a quantitative comparison of contractile dynamics across the two vessel types and cell models that can form the building blocks for multiscale modeling approaches that will link cellular mechanisms to CBF control at the macroscale in health and disease.

## Methods

### Quantification of transmembrane current densities in SMCs and PCs

We utilize patch-clamp electrophysiology data to characterize transmembrane currents in freshly isolated arteriole SMCs (Rubart *et al*., 1996; Filosa *et al*., 2006; Taylor *et al*., 2023) and capillary PCs (Sancho *et al*., 2022, 2024; Klug *et al*., 2023). Examined PCs were isolated by mechanical disruption of a small piece of brain somatosensory cortex or mouse retinas yielding a mixture of proximal and distal PCs. Main ion channels characterized in central nervous system (CNS) capillary PCs include ATP-sensitive (K_ATP_, composed of Kir6.1/SUR2B subunits), inward rectifying (Kir2), voltage-dependent (Kv1), large-conductance Ca^2+^-activated (BK_Ca_) K^+^ channels, and voltage-gated (VDCC) Ca^2+^ channels. Standard electrophysiological equations fit available current-voltage data to independently quantify permeabilities/conductances of these channels, allowing a direct comparison between the two cell types. *Voltage-gated* Ca^2+^ *channels current (I*_*VOCC*_). Whole-cell electrophysiological recordings in cerebral mural cells indicate high density of L-type VOCCs (Anon, n.d.; Rubart *et al*., 1996; Abd El-Rahman *et al*., 2013; Hariharan *et al*., 2020). I_VOCC_ is modeled as a Goldman-Hodgkin-Katz (GHK) current with a Boltzmann-type voltage-dependent activation term (Eqs. A.6–8) (Yang *et al*., 2003; Kapela *et al*., 2008). Whole cell Ca^2+^ permeabilities are determined by fitting voltage-clamp data in freshly isolated cerebral arteriole SMCs (Rubart *et al*., 1996). Recent data from retinal and brain PCs suggest a significantly reduced (i.e. ∼50%) L-type current density relative to arteriole SMCs (Klug *et al*., 2023).

#### Voltage-dependent K^+^ channel current (I_Kv_)

Voltage-gated K^+^ (K_V_) channels, including the K_V_1.2 and K_V_1.5 subtypes, are functionally active in cerebral mural cells. These channels stabilize V_m_ and regulate Ca^2+^ influx and tone in both arteriolar SMCs and capillary PCs (Straub *et al*., 2009; Dabertrand *et al*., 2015; Sancho *et al*., 2024). An effective total I_Kv_ was modeled using a voltage-dependent steady-state activation function described by a Boltzmann equation (Eqs. A.9&10). Whole-cell conductance and V_m_ dependence was determined by fitting data from isolated cerebral capillary PCs (Sancho *et al*., 2024) and SMCs (Dabertrand *et al*., 2015).

#### Large conductance Ca^2+^ -activated K^+^ current (I_BKCa_)

BK_Ca_ channels mediate significant outward K^+^ currents in cerebral mural cells (Dabertrand *et al*., 2012; Hariharan *et al*., 2020). Whole-cell patch-clamp recordings confirm the functional expression of these channels in freshly isolated SMCs and PCs (Hannah *et al*., 2011; Taylor *et al*., 2023; Sancho *et al*., 2024). I_BKCa_ is modeled using a GHK equation (Eqs. A.15–17) and fitted to whole-cell current–voltage data from (Sancho *et al*., 2024) and (Taylor *et al*., 2023) for capillary PCs and arteriolar SMCs, respectively.

#### Inwardly rectifying K^+^ current (K_ir_)

Kir2 channels have been identified in mural cells based on their characteristic electrophysiological signatures and sensitivity to K_ir_ blockers (Filosa *et al*., 2006; Hariharan *et al*., 2020). I_Kir_ current is modeled as a linear current with a conductance that depends on extracellular K^+^ and has voltage-dependent gating captured by a Boltzmann equation (Moshkforoush *et al*., 2020). Whole-cell conductance was determined by fitting voltage-ramp data (Filosa *et al*., 2006; Sancho *et al*., 2024).

### Modeling V_m_ and Ca^2+^ dynamics in Mural cells

A previously presented SMC model (Kapela *et al*., 2008) is adapted to the cerebral microcirculation by integrating arteriole SMCs or capillary PCs electrophysiological data. Schematic diagram of the mural cell model is presented in Fig. 1. The cell is comprised of three distinct components: the plasma membrane, the cytosol, and an intracellular Ca^2+^ store. The plasma membrane includes kinetic descriptions for channels, pumps, and exchangers, accounting for major transmembrane currents identified in brain capillary PCs or arteriole SMCs. Plasma membrane ion channels considered include large-conductance Ca^2+^-activated (BK_Ca_), ATP-sensitive (K_ATP_), voltage-dependent (K_V_) and inwardly rectifying (K_ir_)) K^+^ channels, non-selective cation channels (NSC), and L-type Voltage-Operated Ca^2+^ channels (VOCC). The model encompasses mathematical descriptions for the Na^+^–K^+^-ATPase (NaK) and plasma membrane Ca^2+^ ATPase (PMCA) pumps, the Na^+^-K^+^-Cl^-^ cotransport (NaKCl), and the Na^+^-Ca^2+^ exchanger (NCX). The intracellular Ca^2+^ store, representing predominantly the sarcoplasmic reticulum (SR), is partitioned to uptake and release compartments (Yang *et al*., 2003; Kapela *et al*., 2008) and includes sarcoplasmic reticulum Ca^2+^-ATPase pumps (SERCA), ryanodine receptor (RyR) or IP3 receptor Ca^2+^ channels (IP3R) and a leak current. A biomechanics module relates cytosolic Ca^2+^ levels to circumferential stress and vessel diameter. Our electrophysiological data suggests that densities/activities of some of these components differ between brain arteriolar SMCs and capillary PCs (Klug *et al*., 2023; Sancho *et al*., 2024). This allows us to construct cell-specific models for the two cell types and explore the physiological relevance of the reported differences.

**Figure 1.**
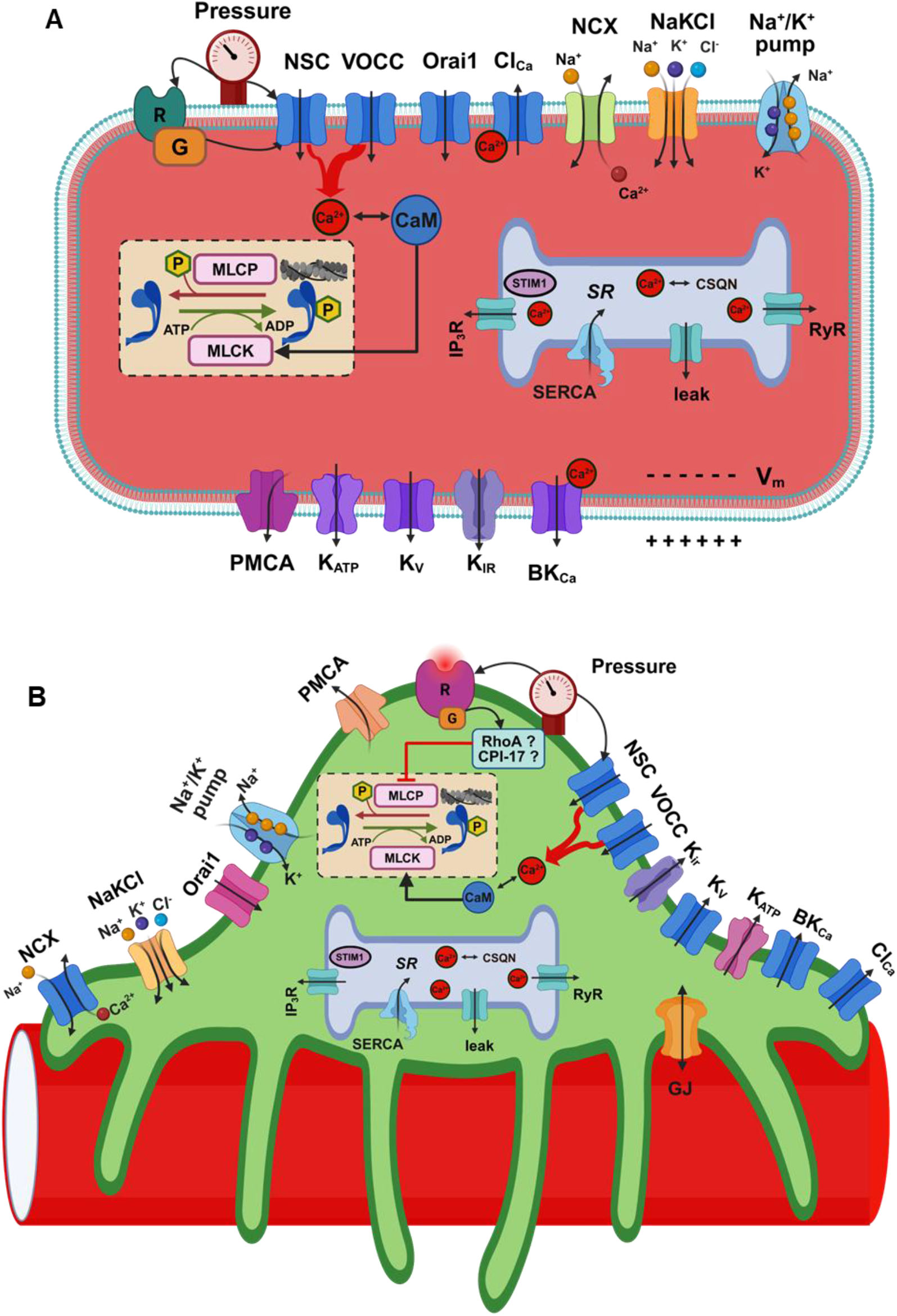
Schematic diagram of model components and their interactions. A mural cell models describes V_m_ and Ca^2+^ dynamics and is adapted to a parenchymal arteriole smooth muscle cell (PA SMC), and a capillary pericyte (PC). The model consider key membrane ion channels, transporters, and receptors, including large conductance Ca^2+^-activated K^+^ channels (BK_Ca_), ATP-sensitive (K_ATP_), voltage-dependent K^+^ channels (K_V_), Inward-rectifying K^+^ channels (K_ir_), Ca^2+^-activated Cl^−^ channels (Cl_Ca_), non-selective cation channels (NSC), store-operated Ca^2+^ channels (Orai1), L-type voltage-operated Ca^2+^ channels (VOCC), Na^+^-K^+^-ATPase (NaK), plasma membrane Ca^2+^-ATPase (PMCA), Na^+^-K^+^-Cl^−^ cotransport (NaKCl), Na^+^-Ca^2+^ exchanger (NCX). Model accounts for intracellular Calcium stores with IP3 (IP3R), and ryanodine (RyR) receptors and sarcoplasmic reticulum Ca^2+^-ATPase pumps (SERCA) and buffering by proteins like Calsequestrin (CSQN) and Calmodulin (CM). Chemomechanical stimuli and G protein coupled receptor signaling can release IP3 and DAG and activate NSC channels.

A detailed description of the modeling approach has been previously presented (Kapela *et al*., 2008) and the reader is referred to earlier publications by us and others for methodological details (Parthimos *et al*., 1999; Yang *et al*., 2003; Tsoukias, 2011). Briefly, membrane electrophysiology is modeled using Hodgkin-Huxley formalism.

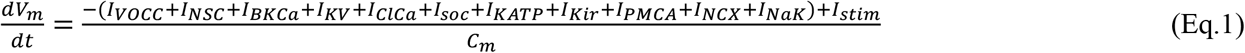

where *V*_*m*_ is the membrane potential, *C*_*m*_ is the membrane capacitance, and *I*_*x*_ represents the current through the respective x ion channel/pump. Mass balances examine the cytosolic dynamics of the four major ionic species (K^+^, Ca^2+^, Cl^-^, Na^+^):

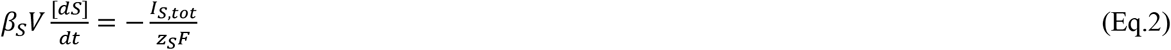

where *V* is the cytosolic volume, [*S*] is the concentration of ionic species *S, I*_*S,tot*_ is the net ionic current entering that compartment, *z*_*S*_ is the ionic valence, and *F* is Faraday’s constant. A fast-buffering approximation is applied to account for the rapid uptake of free Ca^2+^ by cytosolic proteins (i.e. *β*_*Ca*_>1 accounts for reversible binding of Ca^2+^ to intracellular buffers whereas for monovalent ions (K^+^, Cl^-^, and Na^+^, *β*_*S*_ = 1). Similarly, balance equations monitor free Ca^2+^ in the store compartments (Kapela *et al*., 2008).

This detailed approach was successful in capturing salient features of SMC dynamics in our previous work (Kapela *et al*., 2008) and has provided physiological insights into cell and vessel responses to chemomechanical stimuli (Kapela *et al*., 2009, 2010*b*, 2012; Nagaraja *et al*., 2012, 2013; Koide *et al*., 2018). However, the large number of model parameters, many of which cannot be accurately determined from independent experiments, reduces the robustness of previous models. To overcome limitations arising from incomplete characterization of membrane components or uncertainties in quantitation of specific currents, we introduce background (leak) currents (*I*_*bg,K*_, *I*_*bg,Ca*_, *I*_*bg,Cl*_, *I*_*bg,Na*_) that consolidate unaccounted transmembrane ionic fluxes for each ionic species (i.e. K^+^, Ca^2+^, Cl^-^, Na^+^) (Yang *et al*., 2003). For monovalent ions background currents are described using a conductance-based formulation:

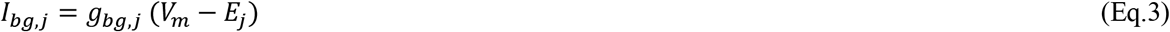

where *g*_*bg,j*_ is the background conductance and *E*_*j*_ is the Nernst potential for ion *j* ∈ {K^+^, Na^+^, Cl^−^}. The background Ca^2+^ flux is expressed using a GHK current formulation, to accommodate the four orders of magnitude transmembrane concentration difference, in line with descriptions often utilized to account for Ca^2+^ currents through different ion channels (i.e. NSC, VOCC):

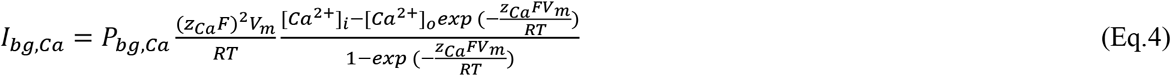

where *P*_*bg,Ca*_ is the permeability, *z*_*Ca*_ the valence, and [*Ca*^2+^]_*o*_ and [*Ca*^2+^]_*i*_ are the extracellular and intracellular Ca^2+^ concentrations, respectively. *R* is the universal gas constant, and *T* is the absolute temperature. Importantly, the conductances/permeability of the background currents are determined by balancing total transmembrane and store currents at rest, given a cell’s resting intracellular ionic concentrations and V_m_. The approach constrains the model to physiological resting values.

Non-Selective Cation channels (NSCs) are expressed in both cerebral arteriolar SMCs and capillary PCs, where they regulate ionic conductance and contribute to pressure- and receptor-induced signaling. A broad range of TRP channels—including TRPC1, TRPC3, TRPC4, TRPC6, TRPV2, TRPV4, TRPM4, TRPM7, TRPP1, TRPP3, and Piezo1—have been implicated in these mural cell types (Tykocki *et al*., 2017; Hariharan *et al*., 2020). These channels differ substantially in their Ca^2+^ selectivity, with reported Ca^2+^:Na^+^ permeability ratios 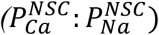 ranging from ∼1 (e.g., TRPC3) to ∼7 (e.g., TRPV4) (Tykocki *et al*., 2017). Such variation implies that the relative distribution of TRP isoforms across mural cells can markedly affect the net 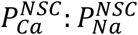. This diversity could provide a mechanistic basis for altered responses between SMCs and PCs (Klug *et al*., 2023; Ferris *et al*., 2025). A net NSC Na^+^ permeability 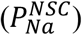 and net 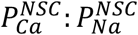 and 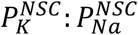 permeability ratios are considered for each cell type, representing weighted average values across the expressed isoforms, and allowing estimation of currents carried by Na^+^, Ca^2+^ and K^+^ ions. Each ionic current is calculated using the GHK current equation:

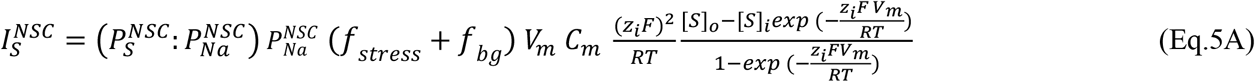

with [*S*]_*o*_ and [*S*]_*i*_ the extracellular and intracellular concentrations of ionic species S, respectively.

Pressure increases affect mural cell tone and yield robust myogenic contractions in microvessels. NSC channels are implicated in pressure-dependent vasoactive responses (Spassova *et al*., 2006; Carlson & Beard, 2011; Hill-Eubanks *et al*., 2014). To account for Ca^2+^ mobilization as transmural pressure increases, a stress-dependent activation of NSC current is assumed (Carlson & Beard, 2011), allowing for stress-induced cell depolarization and Ca^2+^ influx. We assume a sigmoidal dependence of NSC activity on the total circumferential wall stress, *f*_*stress*_ (σ_*tot*_).

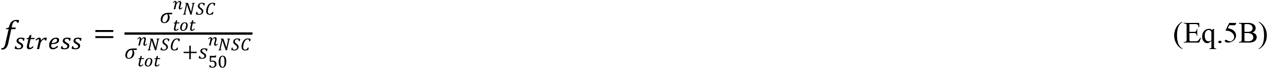

where *n*_*NSC*_ is a Hill coefficient, and *s*_50_ is the stress for half-maximal NSC activation by stress. This formulation links wall stress sensed by mechanosensitive channels or receptors to an effective increase in NSC conductance, to recapitulate the observed relationship between pressure, depolarization, Ca^2+^ influx, and constriction in experimental data.

### Modeling Electromechanical coupling

We utilize the minimal biomechanics model of Carlson and coworkers (Carlson & Secomb, 2005), to translate changes in SMC/PC Ca^2+^ levels to changes in tone and vessel diameter. This earlier model was successful in capturing myogenic constriction in larger arterioles (∼50-∼300μm) and is modified in this study for capturing vessel diameter dynamics in smaller arterioles and capillaries at lower pressures. Similarly to (Carlson & Secomb, 2005), the model accounts for the passive, elastic properties of the vascular wall, as well as for active tone. The passive circumferential stress (σ_pass_) increases exponentially with vessel diameter from a reference zero-stress state (Yang *et al*., 2003):

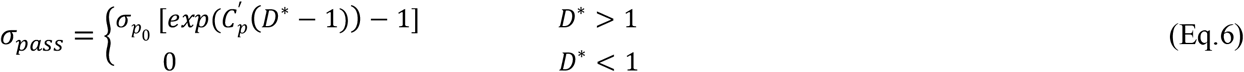

where D* is the vessel’s inner diameter normalized to the diameter at the reference state (*D*_*ref*_). 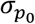 and 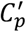 are parameters that determine the exponential increase of *σ*_*pass*_ with D*.

The active circumferential stress depends on the level of SMC/PC tone (i.e. the degree of SMC/PC activation) (*A*_*tone*_), and the stress that the mural cell can generate at a given cell length and saturating Ca^2+^ levels (*σ*_*act*_). *A*_*tone*_ is assumed to have a sigmoidal relationship with [Ca^2+^]_i_.

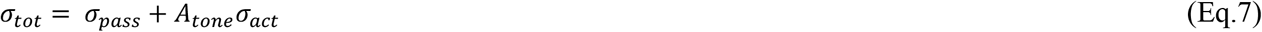

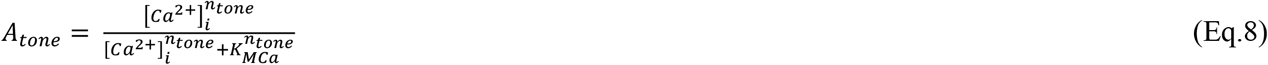

Here, *K*_*MCa*_ is the [*Ca*^2+^]_*i*_ at half-maximal active tone, a measure of the sensitivity of the contractile apparatus to Ca^2+^, and *n*_*tone*_ is the Hill coefficient of the sigmoidal dependence. In previous vascular biomechanics models (Yang *et al*., 2003; Koenigsberger *et al*., 2006; Carlson & Beard, 2011; Kapela & Tsoukias, 2011) the sensitivity of the contractile machinery to Ca^2+^ was assumed constant and independent of mechanical or agonist stimulation. Mural cell myogenic responses may reflect, in part, mechanosensing (i.e. by integrins, Gq_11_ protein coupled receptor (Gq_11_PCRs)), etc.) and the activation of signaling cascades (Rho kinase, CPI-17) that can inhibit MLCP activity and change the sensitivity of the contractile machinery to Ca^2+^, in parallel to any increase in [Ca^2+^]_i_ (Lutz *et al*., 2005; Mederos y Schnitzler *et al*., 2016; Nunes & Webb, 2021). In representative simulations we examined the effect of a pressure-dependent sensitization of the contractile apparatus with slow kinetics (Appendix Fig. A3).

A bell-shaped dependence of active tension at saturating Ca^2+^ (*T*_*act*_*=σ*_*act*_ *w*) on cell’s length was previously assumed (Carlson & Secomb, 2005) and accounts for an optimal cell length for force generation. The description is modified in this study to account for zero active force when the cell’s length decreases below a finite value.

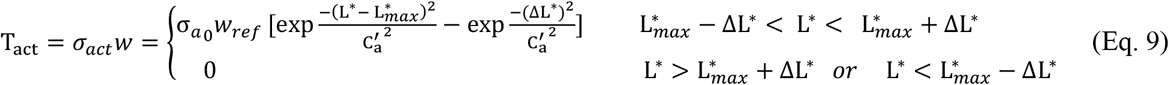

where L^*^ (=L/L_ref_) is the normalized cell length, 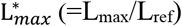 is the normalized cell length for maximum force generation, 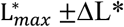 is the normalized length range where the cell can generate contractile force, 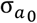 determines the maximum active stress and w, *w*_*ref*_ are the wall thicknesses at the current and reference states, respectively. 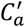 is parameter that controls the width of the bell-shaped dependence of *σ*_*act*_ on L^∗^. In Eq. 9 we assume that a) maximum stress 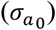 remains effectively constant between vessels of different sizes (i.e. similar actin-myosin density) yielding a linear increase of *T*_*act*_ with vessel size (Carlson & Secomb, 2005) and b) maximum tension 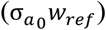 and not stress is preserved when a particular vessel dilates/contracts.

Previous studies often assumed the length of the contracting SMCs to be proportional to the vessel’s inner diameter. Under this assumption, for the inner diameter to approach zero (i.e. as seen in some vessels during maximum constriction), the SMC length needs to approach zero. Furthermore, experimental data suggests a minimum SMC length for force generation typically at 30-40% of the optimal cell length (L_max_) (Mulvany & Warshaw, 1979). Thus, observed changes in inner diameter of contracting microvessels may reflect wall thickening in addition to SMCs shortening. In this study we assume that change in the length of the average SMC reflects a change in the circumference at the midpoint of the vessel wall. During SMC contraction the inner diameter will be reduced by the reduction of the midpoint circumference (*π D*_*mid*_) and the increase in wall thickness (w). Assuming that the cross-sectional area of the vessel wall is preserved during vasoactivity, the following relationship holds between the normalized midpoint 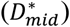 and the inner diameter:

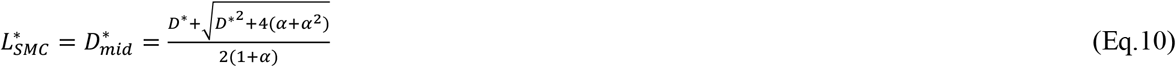

where 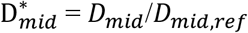 is the midpoint diameter normalized to the reference state and corresponds to the normalized SMC length, 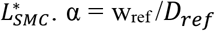 is the ratio of vessel wall thickness to the inner diameter at the reference state. Similarly, we assume that changes in the PC length reflect changes in the outer diameter of the capillary (*D*_*out*_).

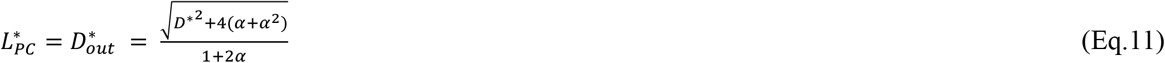

The Laplace law relates total wall stress (σ_tot_), wall thickness (w), and transmural pressure (P) to vessel radius at steady state (Eq. 12):

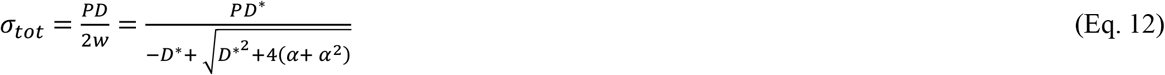

The minimal biomechanical model relates free Ca^2+^ levels in mural cells to the circumferential stress that determines vessel diameter at given transmural pressure. Eq. 5 also relates the activity of NSC channels to circumferential stress, allowing for feedback of transmural pressure on Ca^2+^ dynamics. This allows the model to recapitulate the robust myogenic constrictions in arterioles and capillaries when pressure increases (Nystoriak *et al*., 2011; Dabertrand *et al*., 2015; Klug *et al*., 2023; Ferris *et al*., 2025).

### Quantifying electromechanical responses in pressurized capillaries

We utilize data from two novel ex vivo preparations of CNS microvasculatures (i.e. retinal and cerebral). An isolated en face, intact retina preparation is pressurized after cannulating the ophthalmic artery and allows imaging diameter changes in the arteriole-capillary continuum (Klug *et al*., 2023). Isolated PAs with capillary networks attached are pressurized following cannulation of the PA at one end, tying the other end and sealing the open capillary ends with a second pipette (Ferris *et al*., 2025). Average active and passive (0 [Ca^2+^]_o_) sustained capillary diameter responses during step increases in pressure are normalized to the passive diameter at 5 mmHg (zero stress reference state). Changes in [Ca^2+^]_i_ with pressure were assessed by fluorescence microscopy in transgenic mice expressing the genetically encoded Ca^2+^ indicator GCaMP6f, specifically in mural cells by the *Smmhc*-Cre (ER^T2^) promoter and changes in membrane potential by intracellular electrodes (Klug *et al*., 2023; Ferris *et al*., 2025).

## Results

### Characterization of dominant PC currents

Fig. 2 depicts data from whole-cell patch clamp experiments in freshly isolated mouse cerebral capillary PCs (green circles). Corresponding current-voltage data from arterial SMCs (red circles) (Rubart *et al*., 1996; Filosa *et al*., 2006; Dabertrand *et al*., 2015; Taylor *et al*., 2023; Sancho *et al*., 2024) are also presented and fitted by model equations (lines). Fitting of whole cell voltage dependent Ca^2+^ currents in SMCs (Rubart *et al*., 1996) (Fig. 2A) yields a VOCC Ca^2+^ permeability of 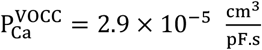. VOCC current densities in retinal and cerebral capillary PC were approximately half of that in arteriole SMCs (Klug *et al*., 2023). Thus a 50% reduction in VOCC is assumed in the PC model. K_V_ (Fiq. 2B) and BK_Ca_ (Fig. 2C) currents in PCs also show a significant reduction (30% and 50%, respectively) compared to SMCs. Kir2 currents at 60 mM extracellular K^+^ (Fig. 2D, inset) were fitted by Eq. S.11 to estimate channel’s conductance 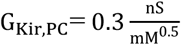. This value is approximately 70% more than the conductance estimated from fitting SMC data, 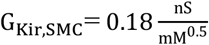 (Fig. 2D).

**Figure 2.**
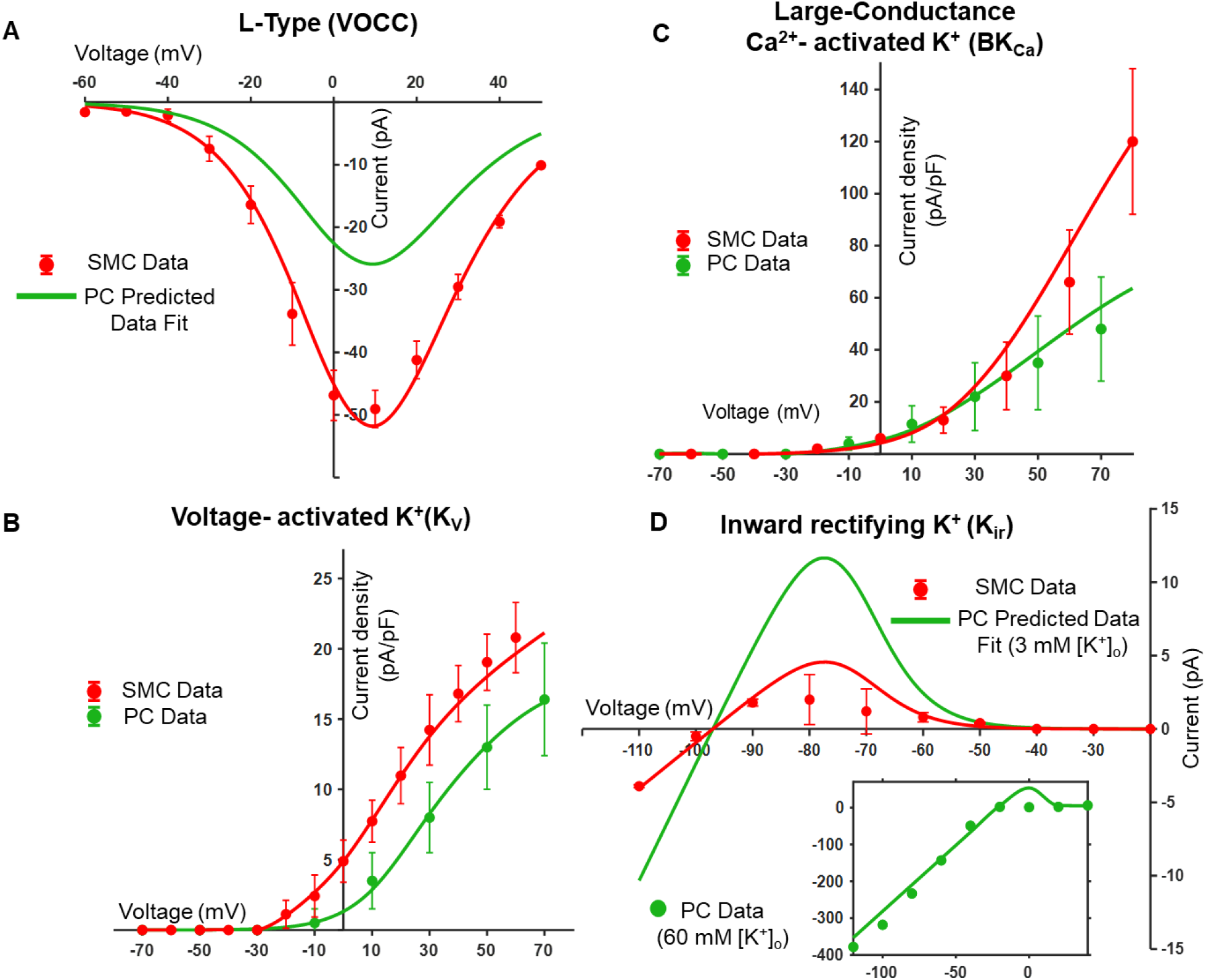
Quantifying current densities of major ion channels in cerebral arteriolar SMCs and brain capillary PCs. (A) L-type Ca^2+^ channels (VOCC). Patch data in SMCs from (Rubart *et al*., 1996) (red circles) are fitted with Eqs. A.6–8 and a permeability of 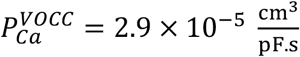 (red line). Predicted VOCC current in PCs assuming a 50% reduction in current density relative to SMCs (Klug *et al*., 2023) (green line, 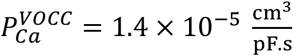). (B) Voltage-activated K^+^ channels (K_V_). Current voltage data in PCs (Sancho *et al*., 2024) (green circles) and in SMCs (Dabertrand *et al*., 2015) (red circles) are fitted with Eqs. A.9&10. Estimated whole-cell conductance 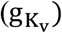 was 1.1 nS for PCs and 1.6 nS for SMCs. (C) Large-conductance Ca^2+^-activated K^+^ channels (BK_Ca_). PCs data from (Sancho *et al*., 2024) (green circles) and SMCs data from (Taylor *et al*., 2023) (red circles) are fitted by Eqs. A.15–17. Estimated whole cell permeability (P_BKCa_) was 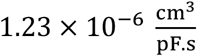 for PCs and 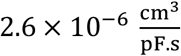 for SMCs. (D) Inward-rectifying K^+^ channels (K_ir_). Whole cell current in SMCs at 3 mM [K^+^]_o_ from (Filosa *et al*., 2006) (red circles) and PCs currents at 60 mM [K^+^]_o_ (green circles, inset) are fitted with Eqs. A.11–13. Estimated conductance (G_Kir_) was 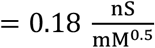 for SMCs and 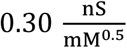 for PCs. Predicted PCs Kir current at physiological [K^+^]_o_ (green line) is almost double the SMCs current (red line). Error bars represent SEM.

### Passive and active stress – strain relationships

Fig. 3 illustrates the predicted biomechanical behavior of an arteriole by the electromechanical model. The blue curve depicts the exponential increase in passive stress (i.e. zero extracellular Ca^2+^) with D*. Active stress, at saturating [Ca^2+^]_i_ (red line) has a bell-shaped dependence on cell length. At the optimum length 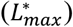 the cell can generate maximum stress that is reduced to zero as the cell length reduces to 50% of L_max_. The maximum stress falls within the range of previous estimates (Carlson & Secomb, 2005). Eq. 10 relates the normalized cell length to the normalized inner diameter that is depicted as a secondary x axis. The total wall stress (black lines) is computed as the sum of passive and active stress components at different [Ca^2+^]_i_. The green line represents Laplace’s law (Eq. 12) and relates total circumferential stress to vessel diameter and transmural pressure. The intersection between the total stress curve at given [Ca^2+^]_i_ and the Laplace law curve defines the equilibrium diameter. As pressure increases, this intersection shifts, as a result of a steeper Laplace curve and an increase in [Ca^2+^]_i_, capturing the interplay between mechanical loading and active tone regulation.

**Figure 3.**
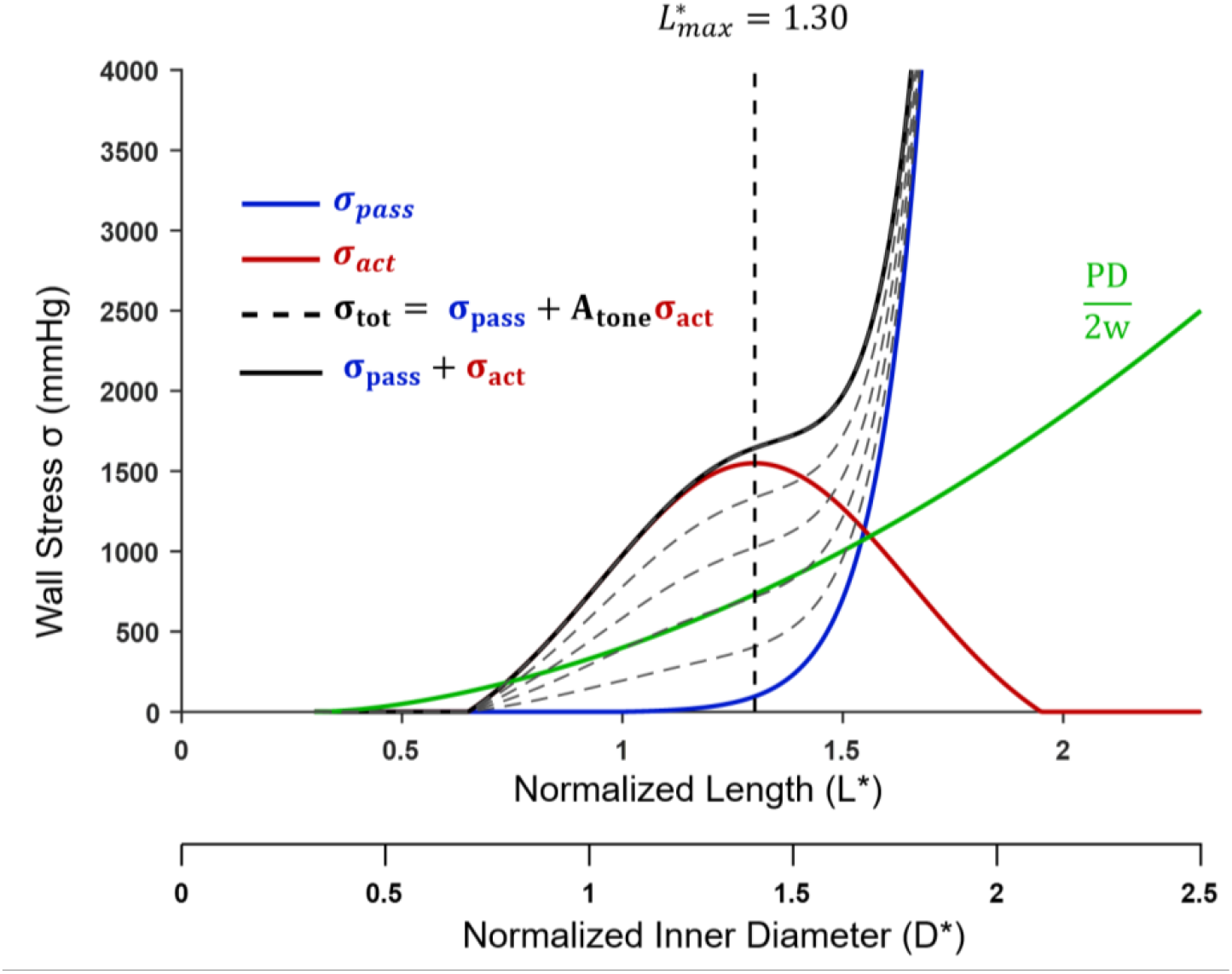
Electromechanical coupling. Predicted stress-strain curves in arterioles. Passive stress (*σ*_*pass*_, blue) and active stress at saturating Ca^2+^ concertation (σ_act_, red) are shown as functions of normalized SMC length (*L*^*\**^) or normalized vessel inner diameter (*D\**). Black lines represent the total wall stress at different levels of SMC activation (*A*_*tone*_) /Ca^2+^ concentration. Total circumferential stress in a vessel at P=40 mmHg transmural pressure (green line) predicted by Lapalace’s law and accounting for changes in wall thickness (w) as diameter (D) changes.

### A model of myogenic reactivity in cerebral SMCs

The SMC model was compared against experimental data characterizing vessel diameter, [Ca^2+^]_i_, and V_m_ in isolated PAs with step increases in intraluminal pressure (Fig. 4A–C). Selected parameters that could not be determined from independent data and with high sensitivity on model outputs (Appendix Fig. A1) were optimized by least-squares fitting. Parameters for the passive stress equation (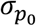 and 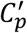), were first determined by fitting the passive dilation data that are independent of electrical or Ca^2+^ dynamics (Fig. 4C; dashed blue line). The NSC’s Ca^2+^ to Na^+^ permeability ratio 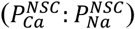 and the maximum PMCA current (*Ī*_*MCA*_) affect Ca^2+^ influx through NSCs and the rate of cytosolic Ca^2+^ sequestration and their values were selected within physiological ranges. Finaly, observed V_m_, Ca^2+^, and diameter responses during myogenic constrictions (Fig. 4) were fitted by optimizing *s*_50_, that determines the sensitivity of NSC channel opening to stress; *K*_*MCa*_, that determines the sensitivity of the contractile apparatus to Ca^2+^; and the active stress parameters (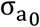 and 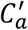).

**Figure 4.**
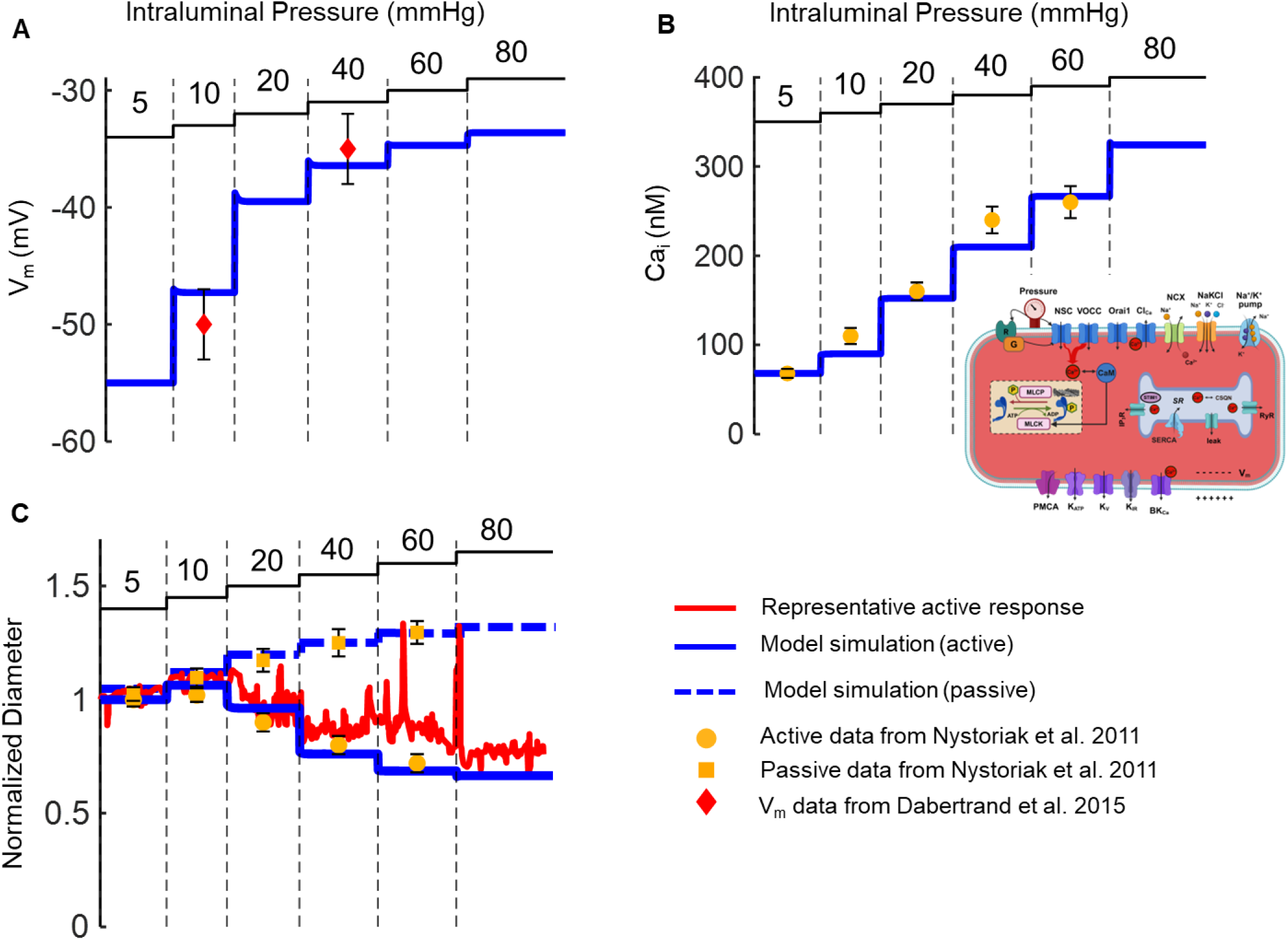
Arteriole response to increases in intraluminal pressure. Changes in V_m_ (A), [Ca^2+^]_i_ (B), and D* (C) during step elevations of transmural pressure. Diameter changes are simulated in the presence (active, solid blue) and absence (passive, dashed blue) of active tone. Simulation results are compared against experimental recordings from (Nystoriak *et al*., 2011; Dabertrand *et al*., 2015) in the presence and absence of extracellular Ca^2+^, respectively.

### Differences in ion channel activities between SMCs and PCs

Recent experimental data by members of our team provided first evidence for robust myogenic constrictions in ACT zone PCs using ex vivo preparation of retinal or cerebral microvasculatures (Klug *et al*., 2023; Ferris *et al*., 2025). Furthermore, we have now characterized main transmembrane currents in freshly isolated brain capillary PCs (Klug *et al*., 2023; Sancho *et al*., 2024). We sought to use this newly acquired data to inform the mural cell model and produce a PC specific implementation. Electrophysiological data suggests differences in densities of key ionic currents between the two cells, including upregulation of K_ir_ activity and downregulation of VOCC, BK_Ca_ and K_V_ activity. We first examined the physiological relevance of these reported differences for the myogenic reactivity of capillaries and tested whether such changes in ion channel activity would allow the robust capillary responses to pressure seen experimentally.

Fig. 5 depicts the optimized SMC model response to pressure (solid red line) from Fig. 4. A 30% reduction in K_V_, a 50% reduction in BK_Ca_ and a 70% increase in K_ir_ current densities did not dramatically change the predicted myogenic response (dotted line). In contrast, subsequently reducing VOCC by 50% significantly compromised the ability of the mural cell model to generate robust constrictions as pressure increases (dashed line). TRPC3, that has relatively low permeability to 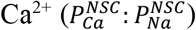 may be a dominant stress activated NSC channel in capillary PCs (Ferris *et al*., 2025). Reducing the NSC net 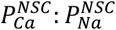 from the control value of 2.5 to 1.6 further reduced the ability of the mural cell to constrict as pressure increases (dashed-dotted line). Experimental data of myogenic constriction in cerebral capillaries from (Ferris *et al*., 2025) are also depicted for reference. They demonstrate more robust myogenic constrictions at lower transmural pressure when compared to responses of arteriolar SMCs. This is in line with the lower operating pressure of capillaries relative to arterioles. Interestingly, the reported differences in ion channel activities in PCs cannot justify the level of myogenic constriction seen experimentally. The significant reduction in VOCC current and/or a decreased Ca^2+^ permeability of stress-activated NSCs would significantly compromise the pressure response if everything else remains unchanged.

**Figure 5.**
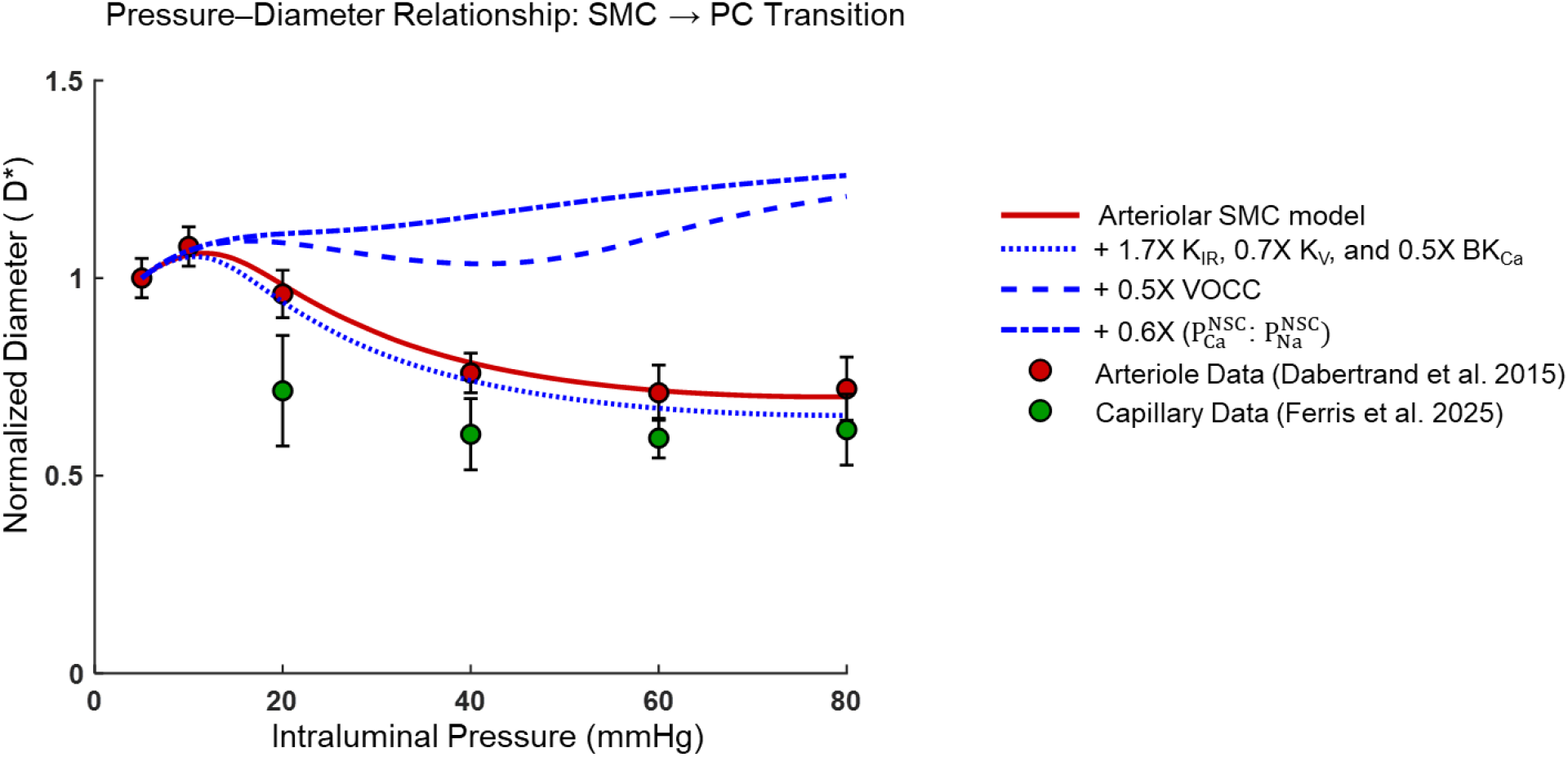
Effect of PC current densities on myogenic tone. Arteriolar SMC model response to step increases in pressure from Fig. 4 (model, red line; and data, red circles). Predicted effect on myogenic tone after sequentially changing current densities of K^+^ channels (K_ir_, K_V_, BK_Ca_) (dotted line), of VOCC (dashed line) and of NSC (i.e. reducing its Ca^2+^ permeability ratio 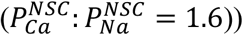, (dash-dotted line). Pressure–diameter data from TZ capillaries (green circles) are included for reference. Measured differences in current densities between PCs and SMCs do not support robust myogenic constrictions in PCs if everything else remains the same.

### A model of myogenic reactivity in capillary PCs

We sought to propose a capillary PCs model, by considering the reported changes in ionic currents and adjusting a minimal number of unidentified parameters, to capture integrated myogenic behavior (Fig. 6). We focused on parameters with high sensitivity on examined cell responses (Appendix Fig. A1&2 and Table A5). The attenuated vasoconstriction predicted after considering PC specific current densities (Fig. 5) can be reversed by reducing the rate of Ca^2+^ sequestration (i.e., reducing the rate of PMCA pump), increasing the sensitivity of the contractile apparatus to Ca^2+^ (i.e. reducing *K*_*MCa*_) and increasing the cell’s mechanosensitivity (i.e. decreasing s_50_) (Table A5). This latter adjustment allowed the PC model to reproduce the earlier onset of depolarization and Ca^2+^ mobilization, allowing for meaningful constrictions at the lower operating pressures of the capillaries. Optimization of passive (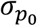 and 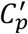) or active (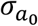 and 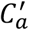) biomechanical parameters produced values that did not significantly differ from corresponding values in the SMC model. Thus, the PC model could recapitulate myogenic constrictions in cerebral or retinal ACT zone capillary PCs (Klug *et al*., 2023; Ferris *et al*., 2025) without assuming significant differences in vascular wall elastic behavior or active force per cross sectional area between PCs and SMCs.

**Figure 6.**
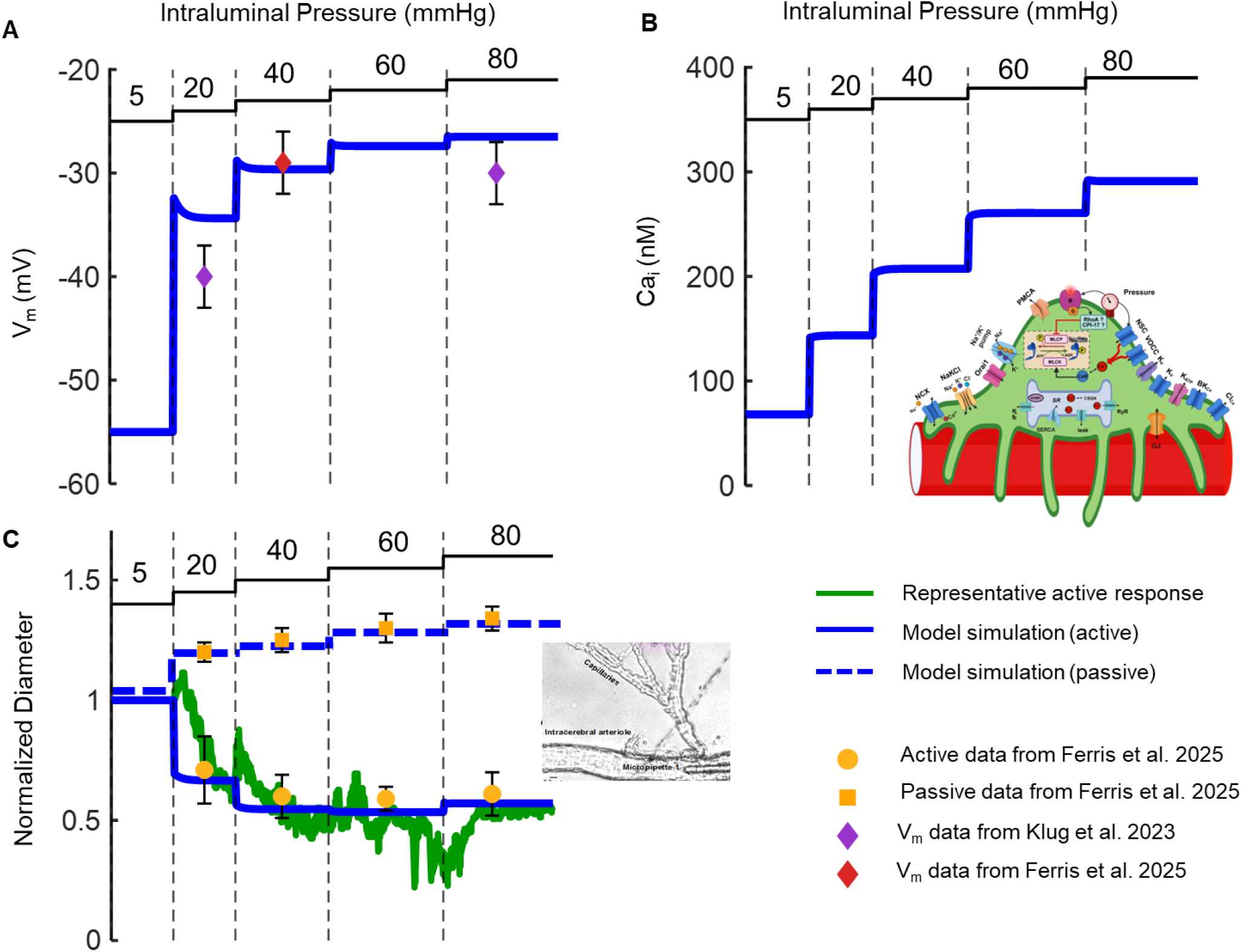
Capillary response to increases in intraluminal pressure. Changes in V_m_ (A), [Ca^2+^]_i_ (B), and D* (C) during sequential step elevations of pressure. Diameter changes are simulated in the presence (active, solid blue) and absence (passive, dashed blue) of active tone. Simulation results are compared against experimental data from (Klug *et al*., 2023; Ferris *et al*., 2025) in the presence and absence of extracellular Ca^2+^, respectively.

### L-type vs NSC Ca^2+^ influx in Myogenic Constrictions

We have recently proposed that the relative contribution of L-type VOCCs to pressure-induced Ca^2+^ mobilization may be progressively reduced as we move down the microcirculation from arterioles to proximal and then distal capillaries (Klug *et al*., 2023). Fig. 7A depicts fluorescent microscopy data from the en face, intact retina preparation in GCaMP6f mice. Changes in [Ca^2+^]_i_ were recorded after pressurizing the ophthalmic artery from an initial value of 20 mmHg to 80 mmHg (red circles/bars). Blockade of the L-type channel with Nifedipine (NIF) shows only partial reduction of the pressure-induced fluorescence increase that could be almost abolished by blockade of TRPC channels with SKF-96365 (SKF) or enhanced by hyperpolarization through K_ATP_ activator Pinacidil (PIN). The latter should create a more favorable electrochemical gradient and enhance Ca^2+^ entry through open NSCs. Thus, data suggests significant VOCC-independent Ca^2+^ influx in retinal capillaries mediated by pressure-induced activation of NSC (TRPC) channels (Klug *et al*., 2023).

**Figure 7.**
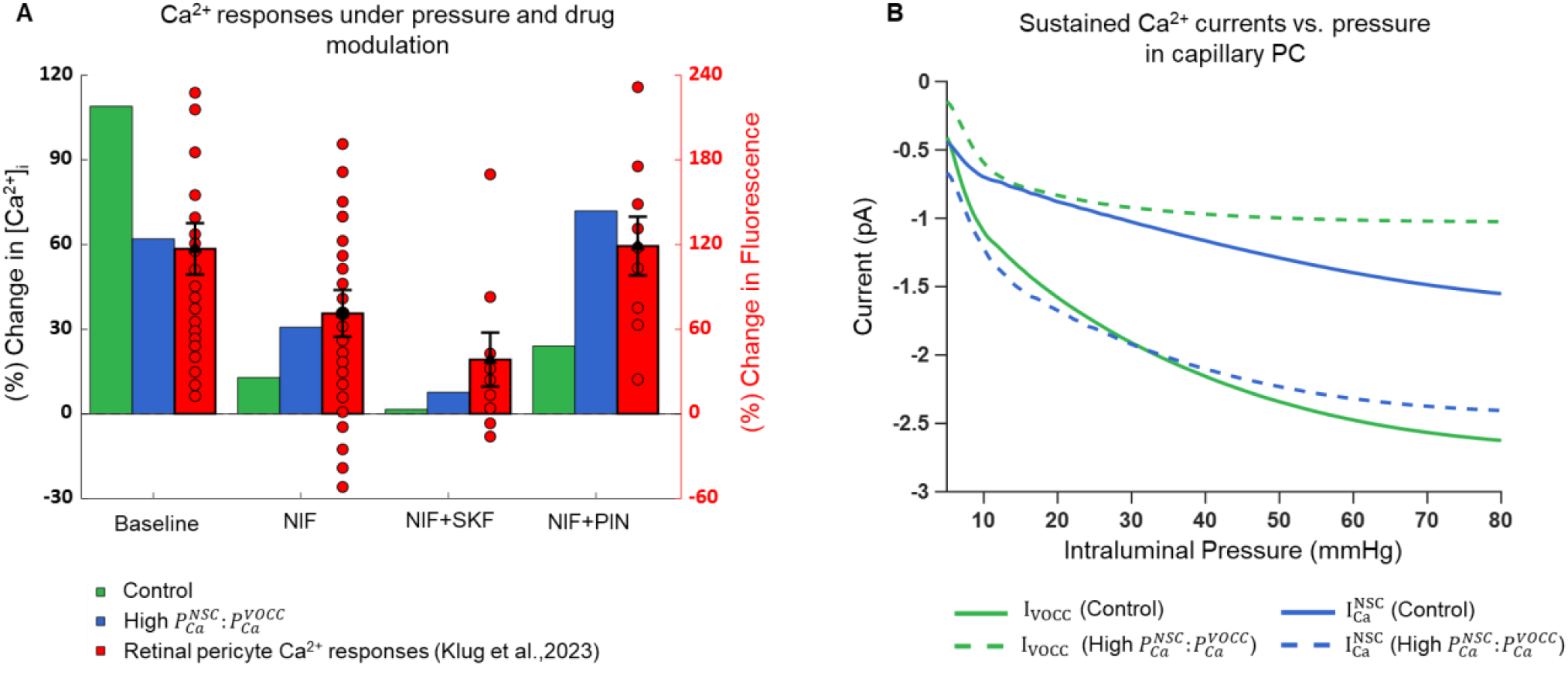
Pressure-evoked Ca^2+^ influx pathways in PCs. (A) Change in Ca^2+^ fluorescence in retinal PCs after increasing perfusion pressure from 20 to 80 mmHg in an isolated intact retinal prep (Klug et al. (2023)) (red circles). Nifedipine (NIF) independent Ca^2+^ influx is sensitive to SKF-96365 (SKF) and Pinacidil (PIN) suggesting significant Ca^2+^ influx through NSC channels. Corresponding model simulations depict change in [Ca^2+^]_i_ with pressure elevation under control conditions (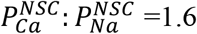 and 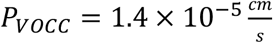) and under a scenario with increased NSC Ca^2+^ selectivity and reduced VOCC permeability (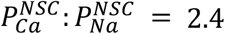 and 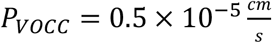). (B) Model-predicted *I*_*VOCC*_ and 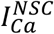 currents as a function of pressure for the control and high 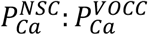 scenarios.

We used the PC model to simulate the experimental protocol. Under control conditions pressure-induced transmembrane Ca^2+^ influx is mostly through the L-type channels and to a lesser degree through NSC channels (Fig. 7B; solid lines). We also examine an NSC favoring scenario by reducing L-type Ca^2+^ permeability by 66% 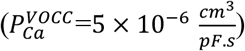 while increasing the NSC Ca^2+^ permeability, 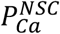, by 50% 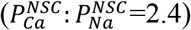, proportionally scaling the currents (Fig. 7B dashed lines). [NSC Na^+^ and K^+^ permeabilities remain constant to provide similar levels of depolarization as pressure increases]. The overall increase of the NSC to L-type Ca^2+^ permeability ratio 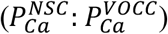 was 4.5-fold. Simulations using the control scenario (green bar) predict a ∼110% rise in intracellular Ca^2+^, as capillary pressure increases from 20 to 80 mmHg, consistent with a robust contraction. The response was largely abolished following blockade of the L-type channels and could not be significantly recovered after hyperpolarization through opening of K_ATP_ channels (i.e. assumed PIN increase in K_ATP_ conductance from 0.03 to 0.3 nS). In contrast the observed trend seen experimentally could be recapitulated using the high 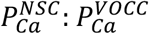 permeabilities ratio scenario (blue bar). Although the baseline Ca^2+^ response was less (∼62%), there was significant VOCC-independent Ca^2+^ increase with pressure that was attenuated with blocking NSCs or increased with hyperpolarization similar to the experiments. Thus, data in retinal PCs suggest a high 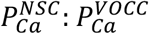 ratio and a significant contribution of NSC Ca^2+^ influx in myogenic constrictions.

### A high 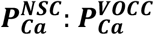 ratio can compromise electromechanical coupling in capillaries

The above data and simulations suggest that influx through VOCC and NSC channels can provide the necessary increase in intracellular Ca^2+^ for robust myogenic constrictions but the relative contribution of the two Ca^2+^ influx pathways in CNS microvascular mural cells may vary as we move from arteries to capillaries. This could potentially alter vasoactive behavior as Ca^2+^ influx through the two channels is affected by electrical signaling in the opposite way. Fig. 8 depicts model response of a pressurized capillary (40 mmHg) upon hyperpolarization (20 pA in every capillary PC). Under control parameter values robust dilation is predicted. Decreasing 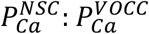 (0.5x) increases the dilatory response; while increasing this ratio (2x) attenuates dilation and makes the capillary insensitive to hyperpolarizing current. For higher 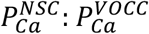 ratios (>2x) constriction in response to hyperpolarizing stimulus is even predicted. Fig. 7B shows the change in transmembrane Ca^2+^ current through VOCC and NCS in each of the above scenarios. Hyperpolarization closes L-type channels, decreasing VOCC-dependent Ca^2+^ influx but increases influx through open NSCs due to the increased Ca^2+^ electrochemical gradient. The net effect for Ca^2+^ flux depends on the relative magnitudes of the changes in the two currents. Thus, a high NSC Ca^2+^ permeability relative to VOCC can decouple the contractile apparatus from electrical signaling or even allow for constriction in response to hyperpolarization.

**Figure 8.**
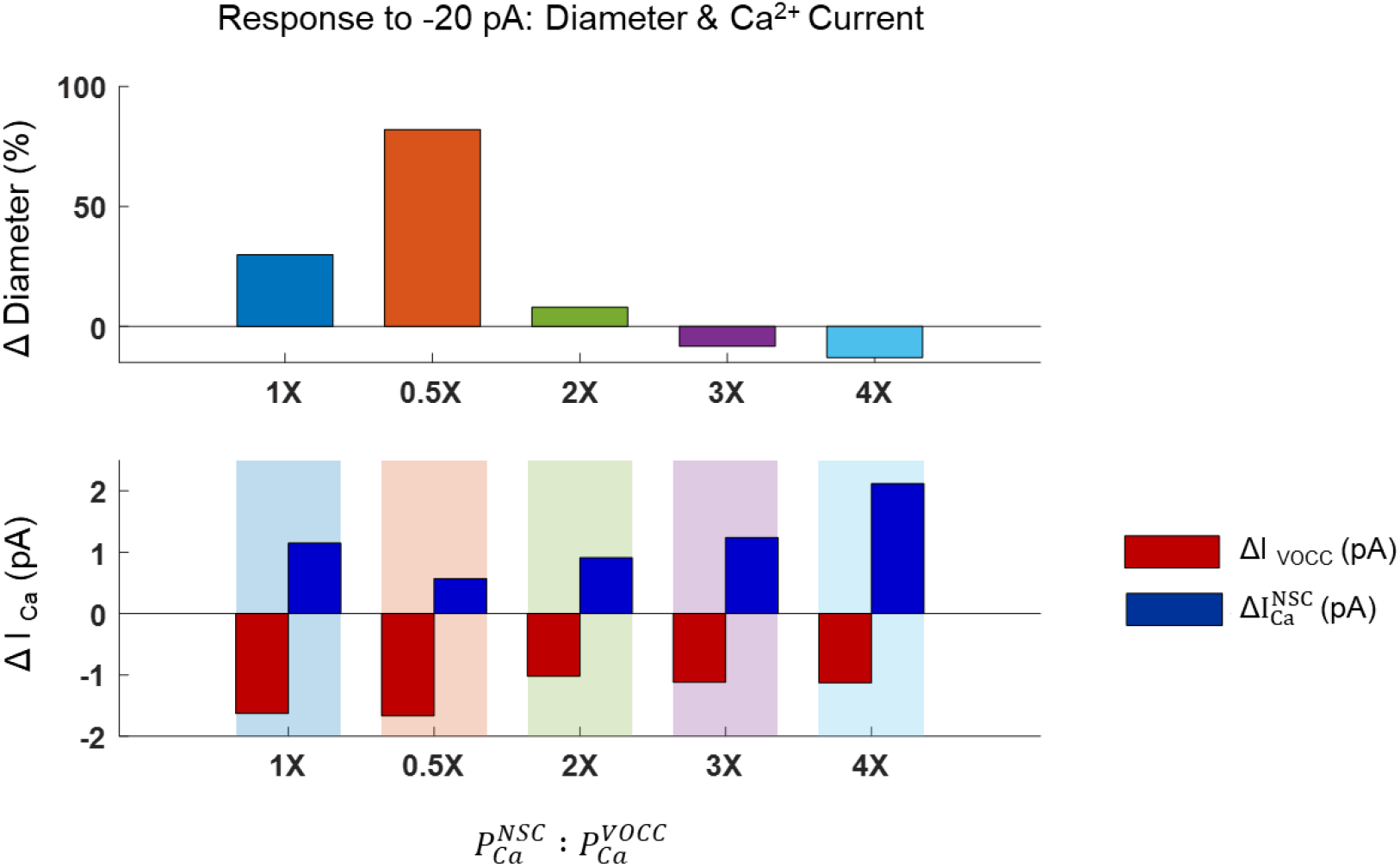
Effect of the NSC-to-VOCC Ca^2+^ permeability ratio on electromechanical coupling. Normalized capillary diameter changes (top) and corresponding Ca^2+^ currents through VOCC and NSC channels (bottom) in response to a 20 pA hyperpolarizing stimulus at 40 mmHg pressure. Simulations were performed across a range of NSC-to-VOCC Ca^2+^ permeability ratio 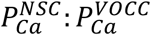 (0.5 to 4 times the control). Increasing the NSC-to-VOCC permeability ratio and shifting the Ca^2+^ influx path from VOCC to NSC, results in a progressive attenuation and eventual reversal of the diameter response. 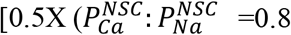 and 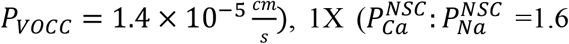 and 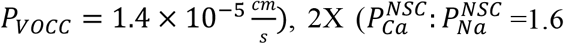 and 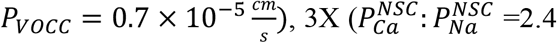 and 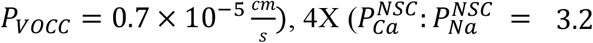 and 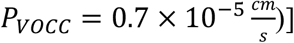.

### Slow PC constrictions by a pressure-induced sensitization of the contractile machinery

Experimental data in capillaries often show slow (i.e. several minutes) constrictions upon stimulation including by step increases in pressure, particularly at low pressures (trace in Fig. 6C) and/or in more distal capillaries (Ferris *et al*., 2025). This behavior cannot be captured with the model where V_m_ dynamics with msec time constants drive Ca^2+^ mobilization in seconds. We examined whether accounting for myogenic constriction through a slow, pressure dependent Ca^2+^ sensitization process can recapitulate experimental observations (Lutz *et al*., 2005; Mederos y Schnitzler *et al*., 2016; Nunes & Webb, 2021; Klug *et al*., 2023; Ferris *et al*., 2025). In a representative simulation (Appendix Fig. A3) we account for a half-maximum Ca^2+^ concentration for constriction (*K*_*MCa*_) that varies from 800 nM to 200 nM as the pressure increases (half-maximum change at P_0.5_= 5 mmHg; t_sens_= 2 min). The modification allows slow diameter responses at low pressure elevations, driven by sensitization of the contractile apparatus being the limiting process in contractile dynamics. Combining a slow Ca^2+^ sensitization with a shift in Ca^2+^ influx from VOCC to NSC allows the model to capture differences between proximal and distal PC dynamics (Appendix Fig. A4).

## Discussion

Recent evidence suggests an important role of capillary PCs in CBF control. PCs can modulate capillary diameter and respond to mechanical forces and neuronal activity, positioning them as important contributors to microcirculatory resistance and neurovascular coupling (Hamilton *et al*., 2010; Hall *et al*., 2014; Hartmann *et al*., 2022). To what extent and under what conditions PC signaling can induce micro- or macroscopic changes, in perfusion is still under investigation (Hill *et al*., 2015; Watson *et al*., 2020; Hartmann *et al*., 2021, 2022) and an integrated view of how capillary and arterial level signaling coordinate appropriate changes in brain tissue perfusion is missing. Here we present a mural cell model that can form the building block for a multiscale modeling approach that will link cell-level signaling to vessel function and brain perfusion at a macroscale. This theoretical framework will allow us to examine how chemomechanical signals along the vascular network intersect to produce appropriate changes in CBF.

We propose a working model of V_m_ and Ca^2+^ dynamics in cerebral SMCs that is informed by electrophysiological recordings in isolated cells. The model is combined with a minimal biomechanics model and can recapitulate tone and diameter responses in cerebral arterioles. This model provides mechanistic insights into how electrical, Ca^2+^ and mechanical signals integrate to regulate arteriolar diameter. The model captures pressure–diameter data in isolated PAs, showing initial vessel distention as pressure increases (<20 mmHg) followed by depolarization via NSC channel opening, voltage-dependent Ca^2+^ entry and constriction at higher pressures. The predicted changes in V_m_ and [Ca^2+^]_i_ closely follow experimental observations, supporting the model’s ability to reproduce physiologically relevant arteriolar behavior across the electrophysiological and biomechanical domains.

PCs in ACT zone capillaries exhibit active tone and robust myogenic constrictions and may play a role as pressure-sensitive regulators of microvascular tone and peripheral resistance. New patch clamp data that characterize currents in freshly isolated PCs and novel ex vivo preparations that allow quantification of proximal capillary responses in the retinal or cerebral microcirculations, enable a first adaptation of the mural cell model into a capillary PC and examination of differences between the two cell types. PCs share a similar ion channel repertoire as SMCs, but a first comparison of major transmembrane currents revealed differences in K^+^ (K_ir_, K_V_, BK_Ca_) and VOCC currents. Importantly, model analysis suggests that a 50% reduction in SMC VOCC density (Klug *et al*., 2023) would have a detrimental impact on their ability to constrict as pressure increases. Accounting for the reported changes in the other examined PC transmembrane currents could not compensate for this effect (Fig. 5). Ex vivo data supports robust myogenic constrictions, and at an earlier onset, as pressure increases in ACT zone capillaries (Fig. 6) that could not be explained by the differences in the measured currents between the two cell types.

Model simulations (Fig. 6) and sensitivity analysis (Appendix Fig. A1 and Table A5) were utilized to identify cell components that can enable sufficient myogenic constrictions at the lower operating pressures of ACT zone capillaries relative to PAs. A reduced rate of Ca^2+^ sequestration (i.e., reducing the rate of SERCA or PMCA pumps or buffering) allows for higher stimulus-induced [Ca^2+^]_i_ increases. However, sufficient sequestration rates are required to allow realistic clearance of Ca^2+^ transients (i.e. following agonist or K^+^ stimuli). Increasing NSC Ca^2+^ influx can compensate for a reduced influx through VOCC but can lead to electromechanical decoupling (see below). Increasing the sensitivity of the contractile apparatus to Ca^2+^ (i.e. reducing *K*_*MCa*_) or increasing the cell’s mechanosensitivity (i.e. decreasing *s*_50_) can also promote myogenic constrictions. Decreasing *s*_50_ was critical for restoring myogenic constrictions in capillaries and capturing their earlier onset relative to the PAs. Slow modulation of K_MCa_, potentially reflecting mechanically-induced signals modulating MLCP activity (Lutz *et al*., 2005; Mederos y Schnitzler *et al*., 2016; Hartmann *et al*., 2021; Nunes & Webb, 2021), enabled the model to recapitulate the observed slow tone development at low pressure elevations in ACT zone capillaries (Klug *et al*., 2023; Ferris *et al*., 2025) (Fig.A3).

We evaluated experimental evidence suggesting an increased importance of Ca^2+^ influx through NSC channels in myogenic constriction of capillaries relative to arteries (Klug *et al*., 2023). Experimental data supports a substantial Ca^2+^ influx through PC NSC in pressurized retinal capillaries. Although this secondary Ca^2+^ influx may compensate for reduced VOCC densities, model simulations highlight a significant functional effect. VOCC and NSC Ca^2+^ influx pathways respond to changes in V_m_ in opposite ways. Depolarization opens VOCC channels but reduces the Ca^2+^ electrochemical gradient. This can result in increasing VOCC Ca^2+^ influx but reducing influx through NSCs. Most importantly, as the ratio of the NSC to VOCC Ca^2+^ influx increases, this leads to electromechanical decoupling and PCs that are insensitive to electrical signaling. Experimental data suggests that this ratio increases as we move distally along the capillary network (Klug *et al*., 2023). The physiological relevance of a spatial gradient in the sensitivity of mural cells to electrical stimuli needs to be further evaluated.

The model adaptations for SMCs and PCs, incorporate different electrophysiology and Ca^2+^ dynamics and can reproduce observed myogenic responses without significant changes in passive or active elastic properties of the tissue wall. The model utilizes a minimal biomechanics description that extends the Carlson and Secomb (Carlson & Secomb, 2005) approach to microvessels at low pressures. This low fidelity approach is ideally suited for multicellular representation of vessels and vascular networks containing large number of mural cells. The proposed framework will allow integrating new experimental data when they become available and explore differences between mural cells from different vessel types of disease conditions.

Mural cell contractile dynamics change as we move from cerebral arterioles to proximal to distal capillaries. Progressively slower responses to vasoactive stimuli (i.e. from a few seconds in arteries to several minutes in distal capillaries (Hartmann *et al*., 2021; Klug *et al*., 2023) may reflect Rho kinase dependent signaling(Hartmann *et al*., 2021). Data also suggest progressively slower myogenic constrictions to pressure steps and a shift from VOCC to NSC mediated Ca^2+^ influx as we move from arterioles to distal capillaries (Klug *et al*., 2023). A progressive shift from Vm/VOCC driven vasoactivity in SMCs, to vasoactive responses driven predominantly by signals modulating a slow Ca^2+^ sensitization at distal PCs can be readily incorporated in the model (Appendix C).

### Limitations

The predictive ability of the detailed cell electrophysiology model is reduced by the large number of parameters and the inability for independent characterization of every cell component using cell specific data. This type of vascular cell model can be useful in formulating hypotheses or setting biophysical limits to proposed mechanisms. In this study, we tried to limit the number of optimized parameters and select physiological values or values used in previous studies for model parameters (i.e. Permeability ratios within the range reported for NSC channels; biomechanical parameters within the range of previous estimates etc.). Although selected free parameters could not be uniquely identified by optimization (i.e., different combination of 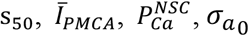 may provide acceptable responses), the model provides appropriate levels of depolarization, Ca^2+^ mobilization and myogenic constrictions to capture observed responses of penetrating arterioles and ACT zone capillaries to transmural pressure elevations. The model can be extended by accounting for more detailed biomechanics including, for example, detailed kinetics of MLCK and MLCP phosphorylation, or the identity of mechanosensors and the mechanisms that transduce chemomechanical stimuli into voltage, Ca^2+^ and diameter responses. Electrophysiological recordings in capillary PCs did not separate between ensheathing PCs in the transitional zone, and thin-strand PCs in distal capillaries. Further experimentation is needed to characterize cell electrophysiology and chemo mechanical signaling with spatial resolution along the capillary space and inform the model.

## Summary

Experimental data and theoretical analysis converge towards a depiction of cerebral mural cells that exhibit significant differences in chemomechanical and electrical signaling along the arteriole to capillary axis (Fig.9). Differences in ion channel activities, in mechanosensitivity and in the Ca^2+^ sensitivity of the contractile apparatus can alter contractile dynamics between arterioles and proximal and distal capillaries. Model simulations examine first evidence on how electrophysiology and vasoactive responses change along the arteriole to capillary continuum. Results suggest that a reduced VOCC density in PCs may compromise contractile dynamics. In ACT zone PCs, any VOCC deficits should be compensated by the upregulation of other cell components to allow for the robust myogenic constrictions documented experimentally. Simulations and data suggest that Ca^2+^ influx through NSC can support tone development but may also decouple the contractile apparatus from electrical signaling. We propose that a higher mechanosensitivity relative to arterioles can allow for robust constrictions at the lower operating pressure of capillaries and that a slow modulation of the Ca^2+^ sensitivity of the contractile apparatus may explain the slower tone development in ACT and distal capillaries. Further experimentation is needed to characterize differences in mural cell dynamics between proximal and distal capillaries and inform the model. Integration of new experimental data into the proposed mathematical framework will provide a platform for assessing electromechanical signaling along the cerebral microcirculation and probe the contributions of distinct ion channels on CBF control and how their dysfunction may lead to perfusion deficits in disease.

**Figure 9.**
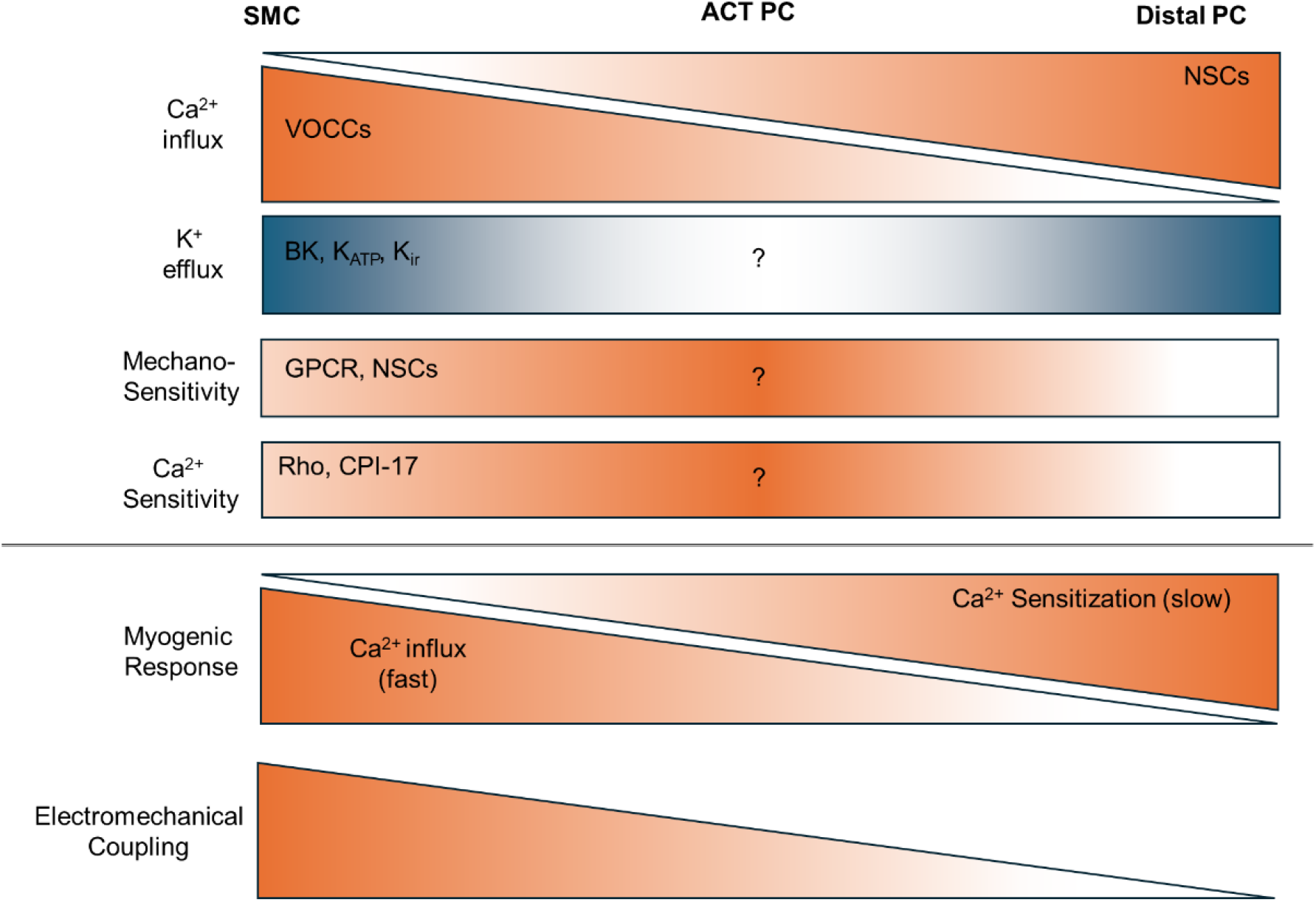
Myogenic tone and electromechanical coupling along the arterial to capillary continuum. Gradients in ion channel activities, mechanosensitivity, and Ca^2+^ sensitivity of the contractile apparatus shape contractile dynamics along the arteriole to capillary axis, with rapid myogenic responses in PAs and progressively slower responses in the ACT zone and distal capillary segments. Experimental evidence supports reduced of VOCC activity in capillary PCs. Ca^2+^ influx through NSC can partially compensate for this reduction, but at the expense of electromechanical coupling, favoring contraction driven by Ca^2+^ sensitization of the contractile machinery with slower kinetics. Further characterization of mural cell electrophysiology and biomechanics across arterioles and proximal-to-distal capillaries can inform the model and improve understanding of contractile dynamics and cerebral blood flow regulation.

## Additional information

### Data availability statement

All data that support the findings of this study are available within the article.

